# Internal Translation of p53 Oncoproteins During Integrated Stress Response Confers Survival Advantage on Cancer Cells

**DOI:** 10.1101/2023.03.03.531004

**Authors:** Maria José López-Iniesta, Rafaela Lacerda, Ana Catarina Ramalho, Shrutee N. Parkar, Ana Marques-Ramos, Bruna Pereira, Lina Miyawaki, Jun Fujita, Roman Hrstka, Luísa Romão, Marco M Candeias

## Abstract

*p53* is the most known and studied tumour suppressor gene. Yet we have recently shown that *p53* is also a proto-oncogene, as it encodes the Δ160p53 oncoprotein. Integrated stress response (ISR) is a survival pathway frequently activated in cancers, marked by the phosphorylation of eukaryotic initiation factor 2alpha (eIF2α) and a defined reprogramming in mRNA translation. Here we identified ISR as a powerful trigger of p53 oncogene, leading to the induction of not only Δ160p53 but also Δ133p53, another protein variant of the *p53* gene. Upon ISR the two isoforms were translated internally from p53 full-length (FL) transcript through an internal regulator of expression site (IRES) located in the vicinity of codon 160. Frameshift mutations upstream of codons 133 and 160 demonstrated that FLp53 protein synthesis is not required for making Δ133p53 and Δ160p53. Instead, targeting IRES(160) with an antisense oligo was sufficient to efficiently and specifically impair the expression of these isoforms without affecting FLp53 levels. This in turn averted ISR’s protective program culminating in cancer cell cycle arrest and death. Mechanistically, FLp53 showed 3 times more affinity to Δ160p53 than to other isoforms, Δ40p53 or Δ133p53. During ISR Δ160p53 localized to the nucleus and strongly inhibited FLp53-mediated activation of pro-apoptotic gene *p53 upregulated modulator of apoptosis* (PUMA). Our results uncover a new branch of the ISR network essential for cancer cell survival and growth and establish the proof of concept for a new strategy to target cancer.

## Δ133p53 and Δ160p53 isoforms are specifically expressed during integrated stress response (ISR)

We have previously shown that the Δ160p53 isoform is naturally pro-oncogenic and provides the survival and invasive advantages observed in cancer cells with a gain-of-function (GOF) mutation in the *p53* gene, R273H^1,2^. Since Δ160p53 is wild-type (wt), naturally occurring and its translation initiation site (TIS) is the most conserved alternative TIS in *p53* among species (**Fig. 1a**), we expected it to play an important physiological role in the cell/organism and also that it might be tightly regulated by cellular/environmental cues. Indeed, we observed induction of Δ160p53 and Δ133p53 in cell lines with endogenous *p53* (human lung carcinoma A549, human fibrosarcoma HT1080, mouse embryonic fibroblast NIH3T3 and human glioblastoma LN229; **Fig. 1b-e**, respectively) as well as in the human lung carcinoma *p53*-null H1299 cell line transfected with Δ133p53 plasmid construct (**Fig. 1f**) or full-length (FL) p53 construct (**Fig. 1g**), when undergoing integrated stress response (ISR); which was achieved through prolonged treatments with endoplasmic reticulum (ER) stressor drugs thapsigargin (Th) and tunicamycin (Tu) and to a smaller degree with DNA damaging agent etoposide (Eto), as confirmed by the increased levels of phosphorylated eIF2α (P-eIF2α), the defining mark of ISR^3^. Mutating the initiation codon for Δ160p53 (M160A) abolished its expression while mutating AUG133 (M133A) increased Δ160p53 levels (**Fig. 1h**), in line with previous reports showing that removal of an upstream TIS leads to enhanced translation from the downstream open reading frame (ORF)^4,5^.

**Figure 1.**
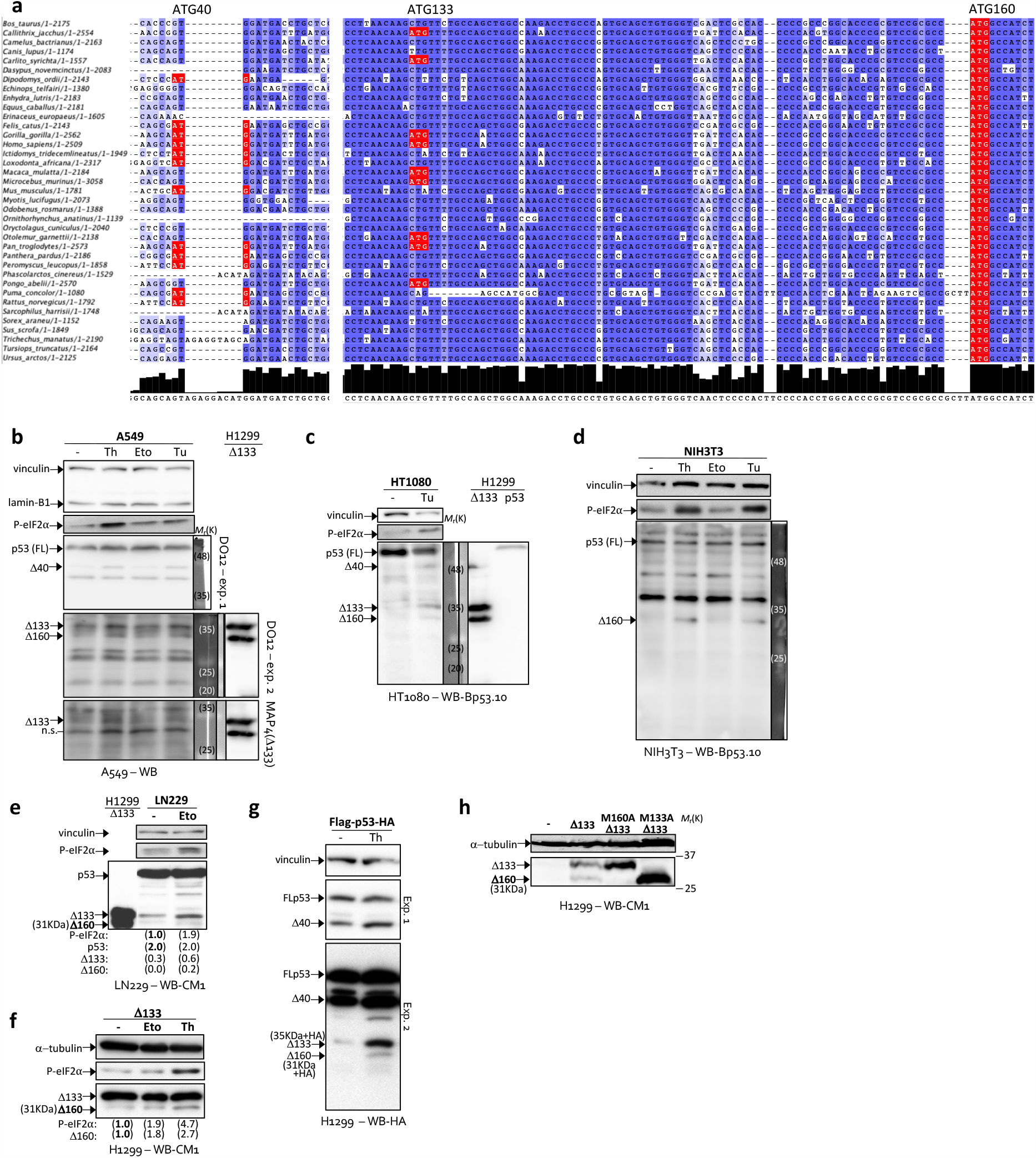
Δ133p53 and Δ160p53 isoforms are specifically expressed during integrated stress response (ISR). **a**, Sequence alignment of regions surrounding internal translation initiation sites (TIS) in *p53* from 37 different mammalian species using MAFFT server within Jalview. TIS are shown in red and were labelled with codon numbers corresponding to the human *p53* sequence: ATG40, ATG133 and ATG160. **b-e**, Western blot analyses (WB) of A549, HT1080, NIH3T3 and LN229 cells endogenously expressing p53 and cultured under normal conditions (-) or integrated stress response (ISR) conditions through treatment with thapsigargin (Th, 16h), tunicamycin (Tu, 16h) or etoposide (Eto, 21h). **f-h**, WB of p53-null H1299 cells expressing no p53 or Δ133p53 wild-type (Δ133) (**f, h**) or tagged full-length p53 (Flag-p53-HA) (**g**) or Δ133p53 mutated at translation initiation starts AUG160 (M160A Δ133) or AUG133 (M133A Δ133) (**h**), and subjected or not to ISR (Th 16h or Eto 36h), as indicated. DO12, monoclonal anti-p53 (aa 256-267) antibody. MAP4, monoclonal anti-Δ133p53 specific antibody. Bp53.10, monoclonal anti-p53 (aa 374-378) antibody. CM1, polyclonal anti-p53 antibody. P-eIF2α, phosphorylated eIF2α (ISR marker). Shown are representative data of at least three independent experiments. The numbers in parenthesis under the WBs specify the amounts of protein for the indicated bands relative to bands showing numbers in bold, according to WB quantifications and normalization against loading control (α-tubulin or vinculin).

## Δ133p53 and Δ160p53 are translated internally from full-length (FL) p53 mRNA during ISR

The fact that Δ160p53 can be stimulated independently of other isoforms, including Δ133p53 at times (**Fig. 1f**), means that Δ160p53 can be regulated post-transcriptionally since Δ160p53 and Δ133p53 are synthesized from the same mature transcripts^2,6^. By using inhibitors of translation (cycloheximide, CHX) or protein degradation (MG132), we could conclude that ISR-dependent induction of Δ160p53 takes place mostly at the translational level since CHX inhibited induction by thapsigargin much more efficiently than MG132 (**Fig. 2a**). TIS160 (AUG160) is actually embedded in an optimal context for translation initiation, even relatively to full-length p53 (FLp53)’s TIS, TIS1, and this was true for most mammalian species tested (**Fig. 2b**). So we investigated if Δ160p53 can be translated internally, since that would enable Δ160p53 to be translated independently of other isoforms^7^. We used a bicistronic construct system^8^ (**Fig. 2c, upper panel**) to investigate internal translation initiation from a Δ160p53 transcript that included 78 nucleotides of its 5’-untranslated region (5’UTR) (up to but not including AUG133). The construct also includes an upstream cistron (EGFP) that is translated *via* canonical 5’ end m7G cap-dependent translation. The downstream cistron (in this case Δ160p53 with the 5’UTR) on the other hand is only translated if there is internal translation initiation^8^. We detected internal expression of Δ160p53 from this construct at levels similar to endogenous expression during ISR (**Fig. 2c, lower panel**). Treating the cells with thapsigargin in this system raised Δ160p53 levels more than 3-fold, similar to what had been observed for endogenous Δ160p53 (**Fig. 2d** and **Fig. 1**). Adding one more codon (ATG133) to the construct allowed us to see that Δ133p53 as well can be translated internally and that now it is under the influence of ISR, as opposed to circumstances when it is located in the vicinity of the 5’-end cap of the mRNA (compare **Fig. 2e** with **Fig. 1f**). In fact we could confirm that in the full-length context (**Fig. 1g**) Δ133p53 is indeed expressed and induced internally by introducing a frameshift mutation in codon 130, upstream of the Kozak sequence for TIS133. The frameshift mutation creates a premature stop codon for all products that start translation initiation upstream of the mutation and these will thus lack the HA-tag at the end of the p53 sequence and not be recognized by the antibody. If the band corresponding to Δ133p53 is indeed due to internal translation then it can escape the effect of the frameshift and be successfully recognized by the anti-HA antibody. In **figure 2f**, fs-130 construct still showed the same Δ133p53 band as the original construct (on the right), while all the bands above Δ133p53 disappeared, proving that the protein of 35 KDa (+HA) is indeed being translated internally and is not a product of protein cleavage. The same could be confirmed for Δ160p53 using a frameshift mutation in codon 157 of *p53* (**Fig. 2f, middle lane** and our previous work^9^).

**Figure 2.**
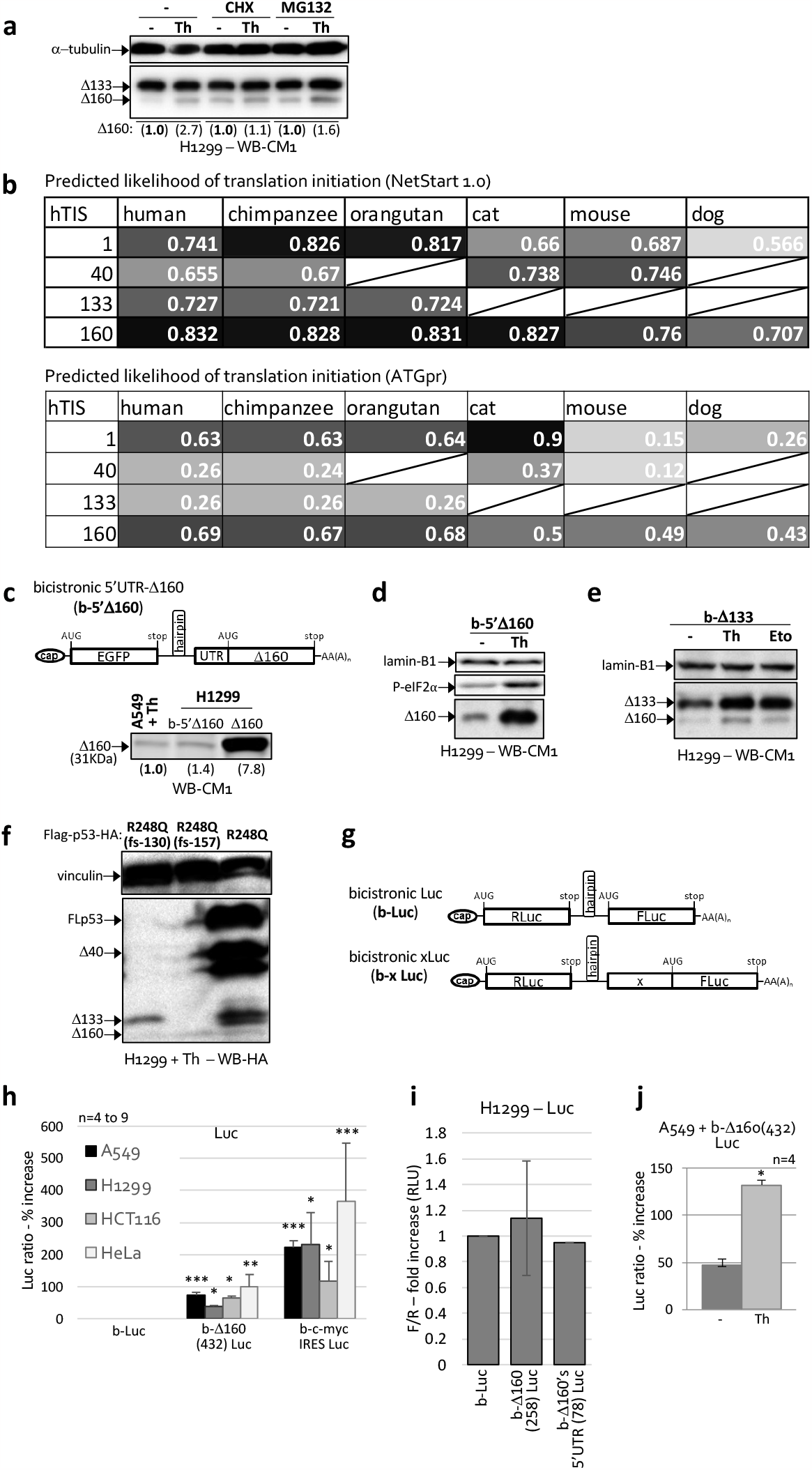
Δ133p53 and Δ160p53 are translated internally from full-length (FL) p53 mRNA during ISR. **a**, Western blot analyses (WB) of H1299 cells expressing Δ133p53 and treated with DMSO (-) or with ISR activator thapsigargin (Th, 16h) and proteasome inhibitor (MG132, 4h) or translation inhibitor (cycloheximide, CHX, 4h). **b**, Likelihood of translation initiation from start codons homologous to human TIS (hTIS) 1, 40, 133 and 160 in 6 different mammalian species, as predicted by NetStart 1.0 (top) and ATGpr (bottom). **c, d**, WB of H1299 cells expressing 5’cap-translation–blocking bicistronic mRNA containing Δ160p53 and its 5’UTR (b-5’Δ160, represented in (**c**), top panel) and treated with DMSO (-) or Th (16h) as indicated. A549 cells endogenously expressing p53 were also analysed for comparison (**c**). P-eIF2α, phosphorylated eIF2α (ISR marker). **e**, WB of H1299 cells expressing bicistronic Δ133p53 mRNA (b-Δ133) and treated with DMSO (-, 36h), Th (16h) or DNA damaging drug etoposide (Eto, 36h). **f**, WB of H1299 cells expressing tagged and mutated p53, as indicated, and treated with Th (16h). fs indicates frameshift mutations of one nucleotide at the indicated codon (157 or 130). **g-j**, Luminescence readings (% increase of Firefly Luciferase / Renilla Luciferase ratio over empty control) of A549, H1299, HCT116 or HeLa cells expressing empty bicistronic dual-luciferase mRNA (b-Luc, represented in (**g**)) or b-Luc mRNAs containing the first 432 or 258 nucleotides of Δ160p53 (b-Δ160(432) Luc or b-Δ160(258) Luc, respectively) or 78 nucleotides of the 5’-UTR of Δ160p53 (b-Δ160’s 5’UTR(78) Luc) or the positive control c-myc IRES (b-c-mycIRES Luc), as indicated, and treated or not with Th (16h). Shown are averages ± s.d. of n experiments as indicated or representative data of at least three independent experiments (\**P* < 0.05, *\*\*P* < 0.01 and *\*\*\*P* < 0.005 compared to negative control). CM1, polyclonal anti-p53 antibody. The numbers in parenthesis under the WBs specify the amounts of protein for the indicated bands relative to bands showing numbers in bold, according to WB quantifications and normalization against α-tubulin.

To further consolidate our finding and also delimitate an internal regulator of expression site (IRES), we performed more studies using different lengths of Δ160p53’s sequence in a dual luciferase reporter system^10^ in four different cell lines (A549, H1299, HCT116 and HeLa), where this time the upstream cistron codes for a Renilla luciferase (RLuc) and the downstream cistron (5’ m7G cap-independent) codes for a firefly luciferase (FLuc) and contains our region of interest as its 5’UTR (**Fig. 2g**). The minimum tested RNA length seen to exhibit IRES activity was 432 nucleotides (nt) starting from AUG160 (**Fig. 2h,i**). As previously observed, ISR enhanced p53 IRES(160) activity by a factor of three (**Fig. 2j**).

Next we wanted to confirm that our observations on IRES activity were not the result of cryptic promoter activity or splicing upstream of FLuc^11^. We first tested for cryptic promoter activity by removing the SV40 promoter from the bicistronic constructs. No cryptic promoter activity was detected except in the positive control, b-MLH1^12^ (**Fig. 3a**). We then used siRNA against the upstream ORF RLuc. This diminished RLuc luminescence and also decreased FLuc signal down to background levels (i.e. b-Luc levels), confirming the integrity of the reporter mRNA molecule and the absence of a cryptic promoter (**Fig. 3b, c**). Lastly, RT-PCRs were performed demonstrating again the existence of a single mRNA form (no splice variants) (**Fig. 3b, d**). This far, our data indicate that Δ133p53 and Δ160p53 isoforms can be regulated *via* IRES-mediated translation during ISR.

**Figure 3.**
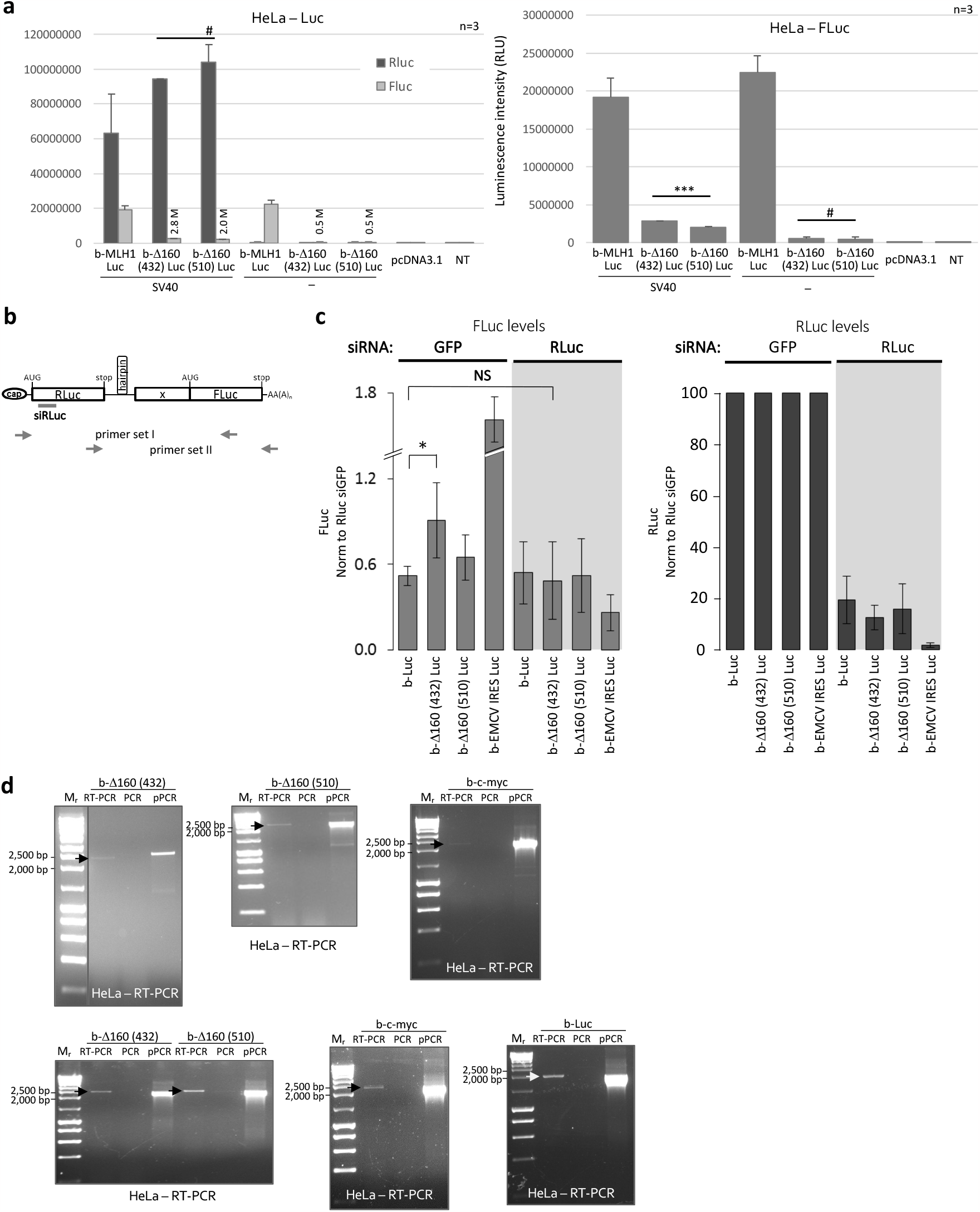
Absence of cryptic promoter activity and alternative splicing in bicistronic dual-luciferase constructs. **a**, Luminescence readings for renilla luciferase (RLuc) and firefly luciferase (FLuc) in HeLa cells not expressing any luciferase (pcDNA3.1 vector and non-transfected (NT)); or transfected with bicistronic dual-luciferase mRNAs containing p53 IRES(160) with 432 nucleotides (SV40 | b-Δ160 (432) Luc), p53 IRES(160) with 510 nucleotides (SV40 | b-Δ160 (510) Luc) or positive control for cryptic promoter activity, MLH1’s 5’UTR (SV40 | b-MLH1 Luc); or transfected with similar bicistronic constructs but lacking the SV40 promoter (labelled “–” instead of “SV40”). **b**, Schematic of the general bicistronic dual-luciferase mRNA (where “x” is replaced with the sequence of study), showing target locations for siRNA used in (c) and PCR primers used in (d). **c**, Luminescence readings for renilla luciferase (RLuc) and firefly luciferase (FLuc) in HeLa cells expressing empty bicistronic dual-luciferase mRNA (b-Luc) or b-Luc mRNAs containing p53 IRES(160) with 432 nucleotides (b-Δ160(432) Luc), p53 IRES(160) with 510 nucleotides (b-Δ160 (510) Luc) or EMCV IRES (b-EMCV IRES Luc), and treated with siRNA control (GFP) or siRNA targeting RLuc, as indicated. **d**, RT-PCR of total RNA extracted from HeLa cells expressing the indicated bicistronic dual-luciferase mRNAs using primer sets I (top panel) or II (lower panel) shown in (c). PCR without RT (PCR) served as negative control and PCR of plasmid DNA (pPCR) as positive control. Shown are averages ± s.d. or representative data of at least three independent experiments (#*P* > 0.05, \**P* < 0.05, and *\*\*\*P* < 0.005 as indicated).

## Δ160p53 acts as a potent and gene-specific transcription regulator co-factor of FLp53

Next we wanted to know how much – and through which mechanisms – the upregulation of Δ133p53 and/or Δ160p53 by ISR can affect cancer cell fitness and tumour growth. We first investigated if these isoforms could act in conjunction with FLp53 since they all contain the oligomerization domain. By placing a Flag tag sequence in the 5’ end of *p53* coding sequence we could produce a construct with the capacity to express the 4 alternative translation isoforms (FLp53, Δ40p53, Δ133p53 and Δ160p53) but where only FLp53 protein carries the Flag tag. To enhance the expression of shorter isoforms we also inserted the R273H mutation^2^. Immunoprecipitation (IP) with an anti-Flag antibody pulled down FLp53 effectively and with it all the detectable amount of Δ160p53; though much less Δ133p53 and Δ40p53 were co-IPed, indicating that Δ160p53 has a much stronger affinity for FLp53 compared to other isoforms (**Fig. 4a**). The same result was observed with the reverse co-IP, i.e., IP of Δ160p53 or Δ133p53 using anti-HA antibody followed by WB to detect untagged wt FLp53 (**Fig. 4b**). These findings, together with the observed translocation of Δ160p53 into the nucleus during ISR (**Fig. 4c**), made us hypothesize that Δ160p53 may be an efficient regulator of FLp53’s trans-activator function. Co-expression experiments showed that Δ160p53 strongly inhibits FLp53-mediated activation of *PUMA*, a pro-apoptotic gene, but had no effect on the expression of *p21* (G1 arrest) or *14-3-3σ* (G2 arrest) (**Fig. 4d**). No changes were observed in mRNA expression in the absence of FLp53 (**Fig. 4e**), confirming that Δ160p53 affects FLp53’s target gene specificity and is unlikely to transactivate genes on its own as it lacks the transactivation domains present in the N-terminus of FLp53.

**Figure 4.**
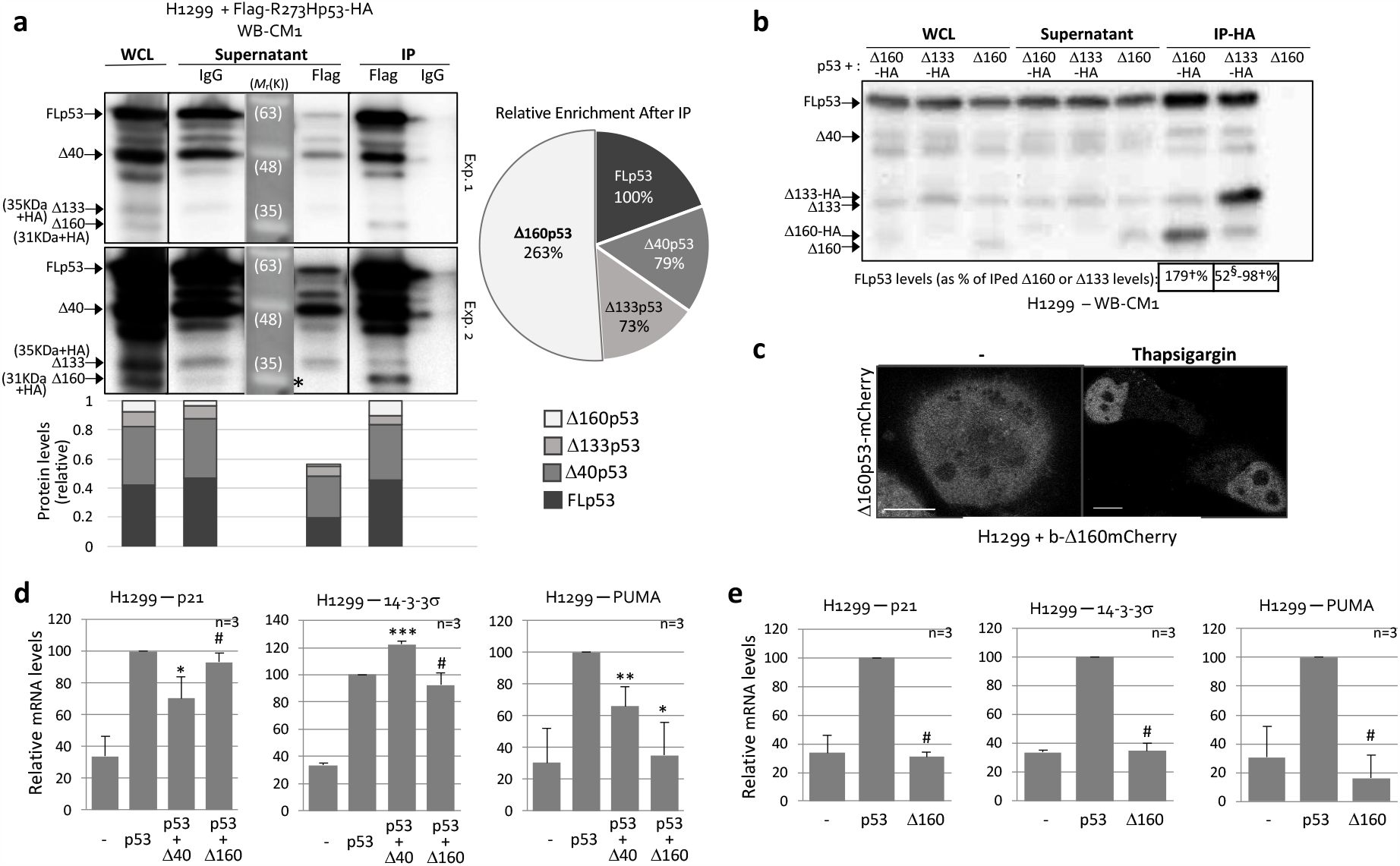
Δ160p53 acts as a potent and gene-specific transcription regulator co-factor of FLp53. **a, b**, Immunoprecipitation (IP) of full-length (FL) FLag-R273Hp53-HA protein with Flag antibody **(a**) or Δ160p53-HA or Δ133p53-HA together with Δ160p53-HA with HA antibody (**b**) followed by Western blot analyses (WB) with polyclonal anti-p53 antibody CM1. WCL, whole cell lysate before IP. Supernatant, supernatant of samples after IP and before washes. IgG, control samples incubated with purified rabbit IgG instead of Flag or HA antibody. The numbers in squares under the WB in (b) specify the relative percentage amount of FLp53 protein compared to the amount of IPed Δ160p53-HA or Δ133p53-HA in the same lane, without any adjustments (†) or considering expression levels of Δ160p53-HA protein from Δ133p53-HA mRNA and equal binding capacity as in the Δ160p53-HA IP sample (§). **c**, Fluorescent imaging of H1299 cells expressing bicistronic Δ160p53-mCherry mRNA and treated with DMSO (-) or ISR-inducing drug Thapsigargin for 16h. **d, e**, RT-qPCR quantifications of indicated endogenous mRNAs from H1299 cells expressing or not different p53 proteins, as specified. Shown are averages ± s.d. of n experiments as indicated or representative data of at least three independent experiments (#*P* > 0.05, \**P* < 0.05, *\*\*P* < 0.01 and *\*\*\*P* < 0.005 compared to p53 alone (d) or no p53 (e)). CM1, polyclonal anti-p53 antibody.

## Targeting p53 IRES(160) specifically inhibits the expression of Δ133p53 and Δ160p53 and impairs ISR-associated cancer cell survival and growth

Following these results we wanted to know if Δ160p53 and p53 IRES(160) have a central role in mediating the survival of cancer cells – since Δ160p53 expression abolished PUMA activation. For this we tested an anti-sense morpholino oligo targeting IRES(160) (MO-IRES or MO-i). MO-IRES efficiently and specifically knocked-down Δ133p53 and Δ160p53, which are both under the control of the same IRES (IRES(160)); but, importantly, unlike siRNA against exons 2-3 that knocked-down all p53 forms including FLp53, Δ40p53 and Δ133p53 (marked with +), MO-IRES did not affect the expression of FLp53 or 40p53, the tumour suppressor isoforms of *p53* (**Fig. 5a-c**). Because MO-IRES only down-regulates Δ133p53 and Δ160p53, i.e. *p53*’s oncoproteins^2,13^, and not FLp53 and Δ40p53, *p53*’s tumour suppressor proteins^14,15^, we expected to be able to inhibit cancer cell survival and growth to *p53*’s fullest capacity when treating cells with MO-IRES during conditions of Δ133p53 and Δ160p53 expression such as ISR. We first tested our hypothesis by expressing Δ160p53 in A549 cells in an IRES-dependent manner using the bicistronic plasmid construct. MO-IRES efficiently induced cell-death and impaired cell-cycle progression (**Fig. 5d, e**). Next we tested the effect of MO-IRES on the function of endogenous p53 proteins. As expected, MO-IRES had no effect under normal growth conditions, when p53 IRES(160) is inactive; however, MO-IRES greatly increased cell death when cells were undergoing ISR (**Fig. 5f**), proving that these internal products of the *p53* gene have pro-survival functions and restrict *p53*’s own tumour suppressive role under conditions of ISR, a common survival program in tumours^16^. We also show that by specifically targeting the internal translation of Δ133p53 and Δ160p53 isoforms and saving FLp53 and Δ40p53 efficiently kills cancer cells. In the future this might become a more effective strategy to treat cancer than to attempt to reactive mutant p53 while leaving Δ160p53 free to inhibit it.

**Figure 5.**
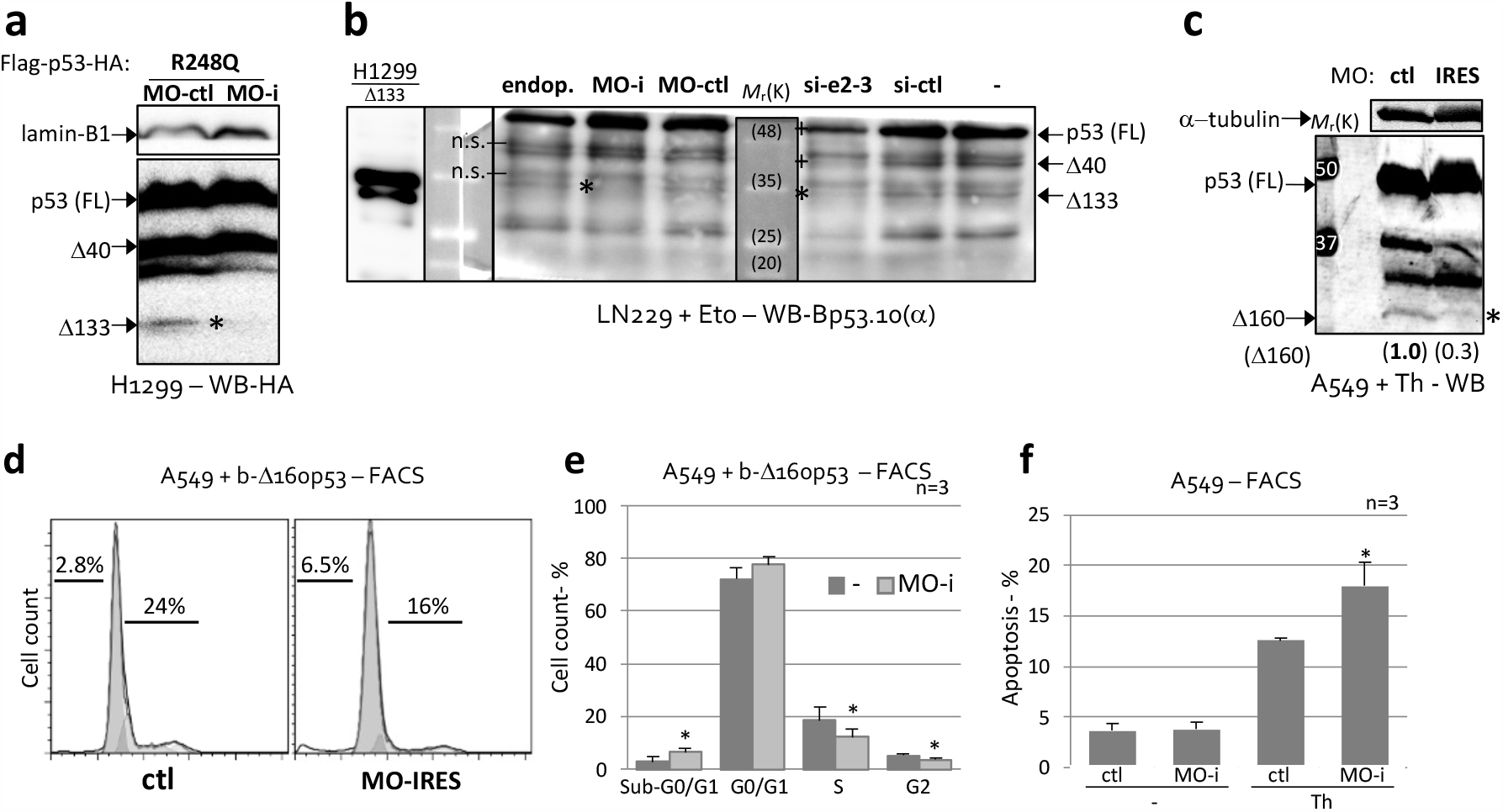
Targeting p53 IRES(160) specifically inhibits the expression of Δ133p53 and Δ160p53 and impairs ISR-associated cancer cell survival and growth. **a**, Western blot analyses (WB) of H1299 cells expressing mutant R248Q p53 and treated with morpholino (MO) against p53 IRES(160) (MO-i) or control MO (MO-ctl). **b, c**, WB of LN299 (**b**) or A549 (**c**) cells expressing endogenous p53 and treated or not with thapsigargin (Th, 16h) or DNA-damaging agent etoposide (Eto, 21h) and morpholino against p53 IRES(160) (MO-i) or control MO (MO-ctl) or siRNA against exons 2 and 3 of p53 mRNA (si-e2-3) or control siRNA (si-ctl) or Endo-Porter alone (endop., MO delivery reagent). * indicates knock-down (KD) of Δ133p53 and Δ160p53; + indicates FLp53 and Δ40p53 KD by siRNA. **d-f**, Fluorescence-activated cell sorting (FACS) analyses of cell-cycle phase and apoptosis in A549 cells expressing endogenous p53 mRNA with (**d, e**) or without (**f**) co-expression of bicistronic Δ160p53 and treated or not with Th (16h) and control morpholino (MO-ctl) or morpholino against p53 IRES(160) (MO-i), as indicated. Shown are averages ± s.d. of n experiments as indicated or representative data of at least three independent experiments (*P < 0.05 compared to control MO). Bp53.10, monoclonal anti-p53 (aa 374-378) antibody. Numbers in parenthesis under the WB specify amounts of Δ160p53 protein for indicated bands relative to band showing number in bold, according to WB quantifications and normalization against α-tubulin.

## Methods

### Cellular assays and reagents

All cell lines were acquired from the American Type Culture Collection (ATCC). Cells were frequently tested for mycoplasma and other contaminations. All the cell lines were maintained at 37°C and 5% CO2 in Dulbecco’s Modified Eagle Media (DMEM) or Roswell Park Memorial Institute (RPMI) media with 10% fetal bovine serum, 5 mM L-glutamine and Pen/Strep 100X solution diluted 1:100. Thapsigargin (Sigma; 0.1-2 μM), Tunicamycin (Sigma; 1.2-2.4 μM) and Etoposide (Sigma; 2.5-3 μM) treatments were for 16-21 hours depending on the cell line and as indicated in the figure legends. Cycloheximide (Cell Signaling; 10 μg/ml) and MG132 (Sigma, Selleckchem; 25 μM) treatments were for 4 hours. For WB, cells were lysed in 1.5x SDS sample buffer (62.5 mM Tris-HCl (pH 6.8), 12% glycerol, 2% SDS, 0.004% Bromo Phenol Blue, and 10% 2-mercaptoethanol). After sonication, proteins were separated by SDS-PAGE on 12% gels and transferred to PVDF membranes (Immobilon-FL, Millipore). After blocking with for 1 h with blocking buffer (1X TBST with 5% w/v non-fat dry milk), membranes were blotted with antibodies diluted in blocking buffer, followed by incubation with secondary antibodies. Membranes were then soaked in Novex ECL (Thermo Fisher Scientific) or SuperSignal West Pico PLUS or ATTO (Thermo Fisher Scientific) and signals were captured with Fujifilm LAS-3000 Imager or imprinted in X-ray films (Amersham Biosciences) using different exposure times and then developed in a Kodak Medical X-ray processor (Carestream Health). For the immunoprecipitation assays, whole cell lysates were pre-cleared with mouse immunoglobulin G and protein G-sepharose before the anti-flag or anti-HA or purified IgG antibody was added. The beads were then washed extensively in buffer A (1% Nonidet P-40, 150mM NaCl, 20mM Tris pH 7.4 in the presence of Complete protease inhibitor cocktail (Roche)) and two times in PBS before boiling in 2x SDS sample buffer. Primary antibodies used for WB were CM-1, Bp53.10, DO12 and MAP4 for p53 isoforms, anti-α-tubulin (Calbiochem DM1A), anti-Flag (Sigma M2), anti-Flag (BioLegend L5), anti-HA (Roche 3F10), anti-Lamin B1 (Santa Cruz Biotechnology A-11), anti-vinculin (Santa Cruz Biotechnology H-10), anti-Phospho-eIF2α (Ser51) (Cell Signaling Technology). Secondary antibodies used were anti-mouse IgG HRP-linked antibody (Cell Signaling Technology), anti-rabbit IgG HRP-linked antibody (Cell Signaling Technology), anti-rat IgG HRP-linked antibody (Cell Signaling Technology), horseradish peroxidase-conjugated goat anti-mouse IgG (BioRad) and goat anti-rabbit IgG (BioRad). All p53 constructs were cloned into pcDNA3.1; dual-luciferase reporter constructs were cloned into psiRF. Morpholino antisense oligos were designed to target the most conserved and stable stem-loop of p53 IRES(160) (SL-V) and block translation of both Δ133p53 and Δ160p53 (MO-IRES or MO-i; Gene Tools, OR, USA). Negative control Morpholino oligos used target a human beta-globin intron mutation that causes beta-thalassemia (MO-ctl or control or -; Gene Tools, OR, USA) or were a mutated version of MO-IRES (MO-ctl or control or -; Gene Tools, OR, USA). 5 μl of MO were added to the cells in 1 ml culturing media followed by 5 μl Endo-Porter delivery reagent and cells were harvested 3 days after.

### Confocal fluorescence imaging

To image Δ160p53-mCherry an IX81 inverted microscope equipped with an FV1000, UPlanSApo 20x 0.75 was used. The confocal aperture size and image size were set at 80μm and at 512 x512 pixels with a zoom factor of 2, respectively. The excitation laser and fluorescence filter settings were as follows: excitation laser 559 nm for mCherry; excitation dichroic mirror, DM405/488/559; mCherry channel PMT dichroic mirror; mCherry channel PMT spectral setting, 570–670 nm.

### Cell-cycle, cell-proliferation & cell-death Analyses

Cells were harvested and fixed with ice-cold 70% ethanol. After 30 min incubation with RNase at 37° C, cells were stained with propidium iodide (PI, Sigma-Aldrich; 50 mg/ml) and cell-cycle distribution was analysed using FACSAria II flow cytometer (BD biosciences) and FlowJo Software^17^. The sub-G0/G1 population represents cells undergoing apoptosis.

### Luciferase assays

Lysis was performed in all cell lines with Passive Lysis Buffer (Promega) and then cells were subjected to a freeze-thaw cycle at -80° C to 37° C and centrifuged at maximum speed for 5 minutes. The cell lysates were used to determine luciferase activity with the Dual-Luciferase Reporter Assay System (Promega) and a GloMax 96 Microplate Luminometer (Promega), according to the manufacturer’s standard protocol. 10 μl of cell lysate were assayed for FLuc and RLuc enzymatic activities. Ratio is the unit of FLuc after normalization to RLuc.

### RNA isolation, RT and PCRs

Total RNAs were extracted from cultured cells using TRIzol reagent (Thermo Fisher Scientific) according to the manufacturer’s instructions. Complementary DNA was prepared using Superscript III reverse transcriptase (Thermo Fisher Scientific) with random hexamer primers according to the manufacturer’s instructions. The relative abundance of transcripts was measured by quantitative PCR using SYBR Green PCR mix (Applied Biosystems). The mRNA levels for each gene were normalized to that of GAPDH and actin. The primers used were as follows (quantitative PCR):

*p21* : 5’-CCTCAAATCGTCCAGCGACCTT-3’ and 5’-CATTGTGGGAGGAGCTGTGAAA-3’

*14-3-30* : 5’-CCAGGCTACTTCTCCCCTC-3’ and 5’-CTGTCCAGTTCTCAGCCACA-3’

*PUMA* : 5’-GACCTCAACGCACAGTA-3’ and 5’-CTAATTGGGCTCCATCT-3’

*GAPDH* : 5’-AAGGTCATCCCTGAGCTGAA-3’ and 5’-CCCCTCTTCAAGGGGTCTAC-3’

*actin* : 5’-TCACCCACACTGTGCCCATCTACGA-3’ and 5’-TGAGGTAGTCAGTCAGGTCCCG-3’

#### Non-quantitative PCR (NZYTech)

*dual luciferase reporter* set I: 5’-GTCTCGAACTTAAGCTGCAG-3’ and 5’-GCAAATCAGGTAGCCCAGG-3’

*dual luciferase reporter* set II: 5’-ATGGCTTCCAAGGTGTACGA-3’ and 5’-ATCGATTTTACCACATTTGTAGAGG-3’

### Sequence analyses

Sequence alignment of *p53* from 37 different mammalian species were performed using MAFFT^18^ server within Jalview. Likelihood of translation initiation from start codons homologous to human TIS 1, 40, 133 and 160 was analysed in 6 different mammalian species using NetStart 1.0^19^ and ATGpr^20,21^.

#### Sequences used are the following

*Bos taurus* (cattle) NM_174201.2

*Callithrix jacchus* (marmoset) XM_002747948.4

*Camelus bactrianus* (camel) XM_010965924.1

*Canis lupus familiaris* (dog) NM_001003210.1

*Carlito syrichta* (tarsier) XM_008062341.2

*Dasypus novemcinctus* (armadillo) XM_012529094.2

*Dipodomys ordii* (kangaroo rat) XM_013013490.1

*Echinops telfairi* (lesser tenrec) XM_013007322.2

*Enhydra lutris kenyoni* (sea otter) XM_022524640.1

*Equus caballus* (horse) XM_023651624.1

*Erinaceus europaeus* (hedghog) XM_007523372.2

*Felis catus* (cat) NM_001009294.1

*Gorilla gorilla* (gorilla) XM_004058511.3

*Homo sapiens* (human) NM_001126112.3

*lctidomys tridecemlineatus* (squirrel) XM_005332819.3

*Loxodonta africana* (elephant) XM_010596586.2

*Macaca mulatta* (Rhesus monkey) NM_001047151.2

*Microcebus murinus* (mouse lemur) XM_012776058.2

*Mus musculus* (mouse) NM_011640.3

*Myotis lucifugus* (bat) XM_006102578.3

*Odobenus rosmarus* (walrus) XM_004398491.1

*Ornithorhynchus anatinus* (Platypus) ENSOANT00000053152.1

*Oryctolagus cuniculus* (rabbit) NM_001082404.1

*Otolemur garnettii* (galago) XM_012806041.2

*Pan troglodytes* (chimpanzee) XM_001172077.5

*Panthera pardus* (leopard) XM_019413568.1

*Peromyscus leucopus* (white-footed mouse) XM_028869449.1

*Phascolarctos cinereus* (koala) XM_020966468.1

*Pongo abelii* (orangutan) XM_002826974.4

*Puma concolor* (puma) XM_025920288.1

*Rattus norvegicus* (rat) NM_030989.3

*Sarcophilus harrisii* (tasmanian devil) XM_031965445.1

*Sorex araneu* (shrew) XM_004604858.1

*Sus scrota* (pig) NM_213824.3

*Trichechus manatus* (manatee) XM_004376021.2

*Tursiops truncatus* (dolphin) XM_019944223.2

*Ursus arctos horribilis* (grizzly) XM_026520889.1

## Acknowledgments

This work was supported mainly by grants PTDC/BIM-ONC/4890/2014 and PTDC/MED-ONC/32048/2017 from the Portuguese Foundation for Science and Technology (Fundação para a Ciência e a Tecnologia; FCT), 18K07229 (KAKENHI) from Japan Society for the Promotion of Science (JSPS), as well as grants/fellowships from Takeda Science Foundation (2017 Medical Research Fellowship (Oncology, Basic)), Astellas Research Foundation for Pathophysiology and Metabolism (project number 203180600044), AXA Research Fund, Ichiro Kanehara Foundation and Kyoto University (Ishizue, year 2021), all attributed to M.M.C and grant UID/MULTI/04046/2013 to BioISI from FCT/MCTES/PIDDAC and by funds from Instituto Nacional de Saúde Doutor Ricardo Jorge. This work was also partially supported by MH CZ - DRO (MMCI, 00209805) and by BBMRI-CZ no. LM2018125 attributed to R.H and by KAKENHI grant JP20K09539 to J.F.. M.J.L.I. and S.N.P. were partially supported by Otsuka Toshimi Scholarship Foundation. A.C.R. is the recipient of a PhD scholarship from the Portuguese Foundation for Science and Technology (2020.06982.BD). p53 antibodies Bp53.10, CM1, DO12 and MAP4 were a gift from Bořivoj Vojtěšek (Masaryk Memorial Cancer Institute, Brno, Czech Republic).

## Author contributions were as follows

M.J.L.I., R.L. and A.C.R. conducted and supervised experiments. L.M., B.P., S.N.P. and A.M.R. conducted experiments. J.F., L.R. and R.H. provided essential reagents and suggestions. M.M.C. conceptualized the study, collected data, conducted experiments, supervised all experiments, interpreted data and wrote the paper. All authors reviewed and were given a chance to comment on the paper.

## Notes

### Competing Interest Statement

The authors have declared no competing interest.

